# Effects of the muscarinic M1 agonist VU0364572 and the M1/M4-prefering agonist xanomeline on cocaine choice in male and female rats

**DOI:** 10.1101/2025.03.03.641295

**Authors:** Samuel A. Marsh, Nicholas Heslep, S. Stevens Negus, Matthew L. Banks

## Abstract

Cocaine use disorder (CUD) is a continuing threat to public health that lacks a Food and Drug Administration approved pharmacotherapy. However, there is evidence to suggest that muscarinic receptor activation may lead to reductions in cocaine taking, acquisition, and choice. Therefore, we use a cocaine-vs-food choice procedure in male and female Sprague-Dawley rats to evaluate the effectiveness of the bitopic M1/M4 preferring muscarinic agonist xanomeline and the M1 muscarinic agonist/positive allosteric modulator VU0364572 (VU’72). We found that repeated xanomeline did significantly attenuate cocaine choice while VU’72 failed to meaningfully alter cocaine choice across subjects. These results suggest that the mechanism by which muscarinic receptors modulate cocaine reinforcement may be mediated either by M4 muscarinic receptors specifically or a combination of both M1 and M4 muscarinic receptors acting in concert. Furthermore, there is evidence that xanomeline may prove a viable pharmacotherapy for treating CUD.

## Introduction

Cocaine use disorder (CUD) remains a public health crisis that affects an estimated 1.4 million individuals according to a 2021 Substance Abuse and Mental Health Services Administration (SAMHSA) study [1]. Furthermore, reports from the National Institute on Drug Abuse (NIDA) indicate that cocaine overdose was the third leading cause of drug overdose-related death in 2022 [2]. These numbers are likely to rise with the advent of the concurrent stimulant crisis occurring the wake of the opioid crisis [3,4]. Yet, there are still no Food and Drug Administration (FDA) approved CUD pharmacotherapy, highlighting the need for preclinical research to identify the neurobiological mechanisms of cocaine reinforcement and evaluate candidate pharmacotherapies in translational animal models.

One neurotransmitter system that is emerging as a potential mediator of cocaine reinforcement is the muscarinic acetylcholine system. Cholinergic projections directly interact with the dopaminergic system, as is shown with cocaine induced increases in acetylcholine levels within the striatum and hippocampus and that were blocked by the introduction of a D1 antagonist [5–7]. Furthermore, there are cholinergic cell bodies projecting to or located within key regions of the mesocorticolimbic dopamine pathway such as the nucleus accumbens (NAc), ventral tegmental area (VTA), and prefrontal cortex (PFC) [8–11]. The M1 and M4 muscarinic acetylcholine receptors, specifically, are highly expressed throughout both the cortex and dopamine reward circuitry [8,12–14]. Consistent with this pattern of muscarinic receptor expression, cocaine-induced NAc dopamine increases were enhanced in M4 knockout mice [15]. With further evidence suggests that M4 receptor activation opposes the effects of NAc D1-like receptor activation[15,16]. Other evidence suggests that M1 receptors express excitatory pyramidal neurons projecting from the PFC to the NAc, opposing D2-like receptor activation. [17,18] In summary, there is extant neurobiological evidence to suggest that M1 and M4 muscarinic ligands modulate dopaminergic transmission in ways that may support their preclinical evaluation as candidate CUD pharmacotherapies.

Consistent with the neurochemical evidence described above, behavioral studies also support a role of M1 and M4 receptors in altering cocaine’s rewarding and reinforcing effects. For example, cocaine-induced conditioned place preference was attenuated in M1 knockout mice compared to wild type [19]. In addition, a single administration of the M1 bitopic agonist VU0364572 (VU’72) attenuated both cocaine self-administration in a cocaine-vs.-food choice procedure and cocaine-induced dopamine increases within the mPFC and the NAc [20]. Furthermore, treatment with the M1/M4 preferring muscarinic agonist xanomeline decreased cocaine self-administration and cocaine discrimination in rodents [21,22]. Overall, these behavioral results support the continued evaluation of the role of M1 and M4 muscarinic receptors in cocaine self-administration endpoints towards the development of muscarinic agonists as candidate CUD pharmacotherapies.

The aim of the present study was to expand upon these previous findings by determining the effectiveness of VU’72 and xanomeline treatment to attenuate cocaine-vs-food choice in male *and* female rats, as previous self-administration studies only used male rodents. Additionally, we endeavored to determine VU’72 treatment effectiveness to alter cocaine self-administration during a repeated and acute dose regimen. We used a cocaine-food choice procedure to determine muscarinic ligand effectiveness because the primary dependent measure, behavioral allocation (i.e. “drug choice), is less sensitive to reinforcement-independent rate-altering effects of pharmacological treatments than conventional single-operant drug self-administration procedures [23]. Lastly, to confirm the efficacy of our experimental procedure to measure differences in cocaine-induced behavioral allocation we used differential reinforcer (i.e. Ensure™ concentration) as a positive control. We hypothesized that VU’72 and xanomeline administration would attenuate cocaine choice in a similar manner to our positive control. Confirmation of our hypothesis would support the continued development and evaluation of these ligands as candidate CUD pharmacotherapies.

## 2.0 Methods

### 2.1 Subjects

A total of 24 (12 Male/ 12 Female) Sprague-Dawley rats weighing approximately 225 – 275g for females and 250-300g for males were purchased from a commercial supplier (Envigo, Fredrick, MD, USA). All subjects were single-housed in a temperature-controlled and AAALAC-accredited vivarium under a 12 h light:dark cycle with the dark cycle running from 6 pm to 6 am. Subjects were given ad libitum access to both food (Tekland Rat Diet, Envigo) and water in their home cage and were weighed weekly. Animal research and maintenance were conducted following the 2011 NIH Guide for the Care and Use of Laboratory Animals, eighth edition. [24] All enrichment and experimental protocols were approved by the Virginia Commonwealth University Institute for Animal Care and Use Committee.

### 2.2 Catheter Implantation and Maintenance

An indwelling jugular intravenous catheter and vascular access port were aseptically implanted as previously described in Townsend 2021 [25]. Catheters were flushed daily with 0.1 ml of cefazolin (50 mg/ml) and heparin (250 U/ml). Catheter patency was confirmed at the end of each experiment by administering intravenous (IV) methohexital (1.6 mg) and observing instant loss of muscle tone and ambulation.

### 2.3 Apparatus

Modular operant chambers (Med Associates, St. Albans, VT) housed in sound-attenuating chambers (Med Associates) were utilized for these cocaine choice studies as described previously [25]. In brief, each retractable lever had tricolor LED lights (red, yellow, green) located directly above the lever. The syringe pump (PHM-100, Med-Associates) was connected to a fluid swivel (375/22PS, Instech Laboratories) and the IV line was protected by a stainless steel, magnetic tether (Instech Laboratories). The operant chamber also contained a retractable dipper (Med Associates) with a 0.1 ml cup for liquid food delivery (vanilla flavored Ensure® in tap water, Abbot Laboratories, Chicago, IL). All behavioral experiments were conducted using custom programs written in Med-State notation (Med Associates).

### 2.4 Cocaine-vs-Food Choice Training

All behavioral sessions were conducted between 8 AM and 11 AM Monday through Friday. Subjects were initially trained to respond for 0.32 mg/kg/infusion cocaine under a fixed-ratio (FR)1 schedule of reinforcement during daily 2-h sessions. Before each session, subjects received a non-contingent 0.32 mg/kg cocaine infusion followed by a 60-s timeout. Once subjects earned ≥ 20 cocaine infusions, the FR value was initially increased to FR3 and then FR5. Following three consecutive days of ≥ 20 cocaine infusions at FR5, food-maintained responding was trained. Initially, the response requirement was FR1 and response requirement completion resulted in presentation of 32% vanilla flavored Ensure diluted in tap water. Training at FR1 continued until >100 reinforcers were earned during the 2-h session. Then the response requirement was increased first to FR3 and then FR5. One subject had the ensure concentration increased to 100% due to failure to meet acquisition criteria. Subsequently, subjects were trained on the terminal cocaine-vs-food choice procedure. The terminal choice procedure had five separate 20-min components with a 5-min timeout between each component. During the timeout, subjects received a non-contingent food presentation and a non-contingent infusion of the cocaine dose available during the subsequent response component as previously described. [25] Increasing cocaine doses (0, 0.32, 0.1, 0.32, 1.0 mg/kg/infusion) were available as the alternative to food during successive components under a concurrent FR5:FR5 schedule of reinforcement. Subjects could complete up to 10 ratio requirements across both levers, and responding on one lever before completing the ratio requirement reset the FR for the other lever. Cocaine choice was considered stable when the smallest cocaine dose that maintained ≥80 percent cocaine choice did not vary by more than 0.5 log units over three consecutive days of testing.

### 2.5 Experimental Manipulations

Firstly, as a comparator to the pharmacological manipulations, we determined the effects of different ensure concentrations on cocaine-vs-food choice. Each ensure concentration (Water, 18, 32, and 100 % ensure) was determined for five consecutive days and tested using a Latin square design. Different ensure concentrations were determined on successive weeks.

Second, we determined the effects of repeated 5-day xanomeline (1.0 – 10 mg/kg, SC) treatment on cocaine-vs-food choice. An experimental timeline is shown in Figure 2A. Xanomeline was administered as a 5-min pretreatment to the daily choice session. Different xanomeline doses were tested on different weeks and were separated by at least one no-treatment week.

Third, we determined the effects of a single saline or VU’72 dose (0.32 – 10 mg/kg, IP) administered as a 30-min pretreatment to the choice session and monitored choice behavior for two. An experimental timeline is shown in Figure 3A. Initially, saline and 1 mg/kg VU’72 were counter-balanced via Latin square. Subsequent VU’72 doses of 3.2, 0.32, and 10 mg/kg were tested consecutively. In addition, after two weeks following each single VU’72 dose administration, repeated 5-day VU’72 treatments were conducted with the same VU’72 single dose as shown in Figure 4A. Thus, each single VU’72 dose, and saline experiment was followed by a subsequent week of repeated 5-day dosing treatment.

### 2.6 Data Analysis

The primary dependent measures were percent cocaine choice, defined as [(the total number of cocaine reinforcers earned/ total number of reinforcers earned)*100] and reinforcers per component defined as the total number of reinforcers earned in each component. These measures were plotted as a function of the unit cocaine dose. Additional dependent measures were the number of reinforcers (total, cocaine, and food) per session. The primary independent variables were VU’72 and Xanomeline dose as well as Ensure concentration. Data from the last two days of each treatment period were analyzed using either a one-way or two-way repeated-measures ANOVA, mixed-effect analysis, or t-test, as appropriate, with cocaine, VU’72 and Xanomeline dose or ensure concentration as the main factors. Post-hoc comparisons following a significant treatment effect and/or interaction were made to vehicle for drug treatments using a Dunnets post-hoc test or pairwise comparisons were made for ensure concentration using a Tukeys post-hoc. The Geisser-Greenhouse correction was used to account for any sphericity violations. Statistical significance was established *a priori* at the 95% confidence level (p<0.05).

### 2.7 Drugs

Cocaine HCl was provided by the National Institute on Drug Abuse Drug Supply Program (Bethesda, MD). VU0364572 HCl was synthesized by the Vanderbilt VICB Molecular Design and Synthesis Center (Vanderbilt University, Nashville, TN). Xanomeline oxalate was purchased from a commercial vendor (Tocris, Minneapolis, MN). All solutions were dissolved in a bacteriostatic saline and IV solutions passed through a 0.22 um sterile filter before use. All drug doses are expressed as the salt forms listed above.

## 3.0 Results

### 3.1 Ensure concentration manipulations on cocaine choice

Figure 1 shows the effects of different ensure concentrations (0% – 100%) on cocaine-vs-food choice in male and female rats. Increasing ensure concentrations resulted in a concentration-dependent decrease in cocaine choice compared to water (cocaine dose: F(2.0, 17.8) = 56.9, p<0.0001; ensure concentration: F (2.1, 18.7) = 22.2, p<0.0001; interaction: F(3.6, 25.7) = 7.1, p = 0.0008; Figure 1A). Increasing ensure concentrations also significantly increased session food reinforcers and decreased session cocaine reinforcers in a concentration-dependent manner without altering session total reinforcers. (ensure concentration: F(1.9, 18.5) = 4.17, p = 0.035; interaction: F(3, 20.7) = 28, p < 0.0001; Figure 1B).

**Figure 1:**
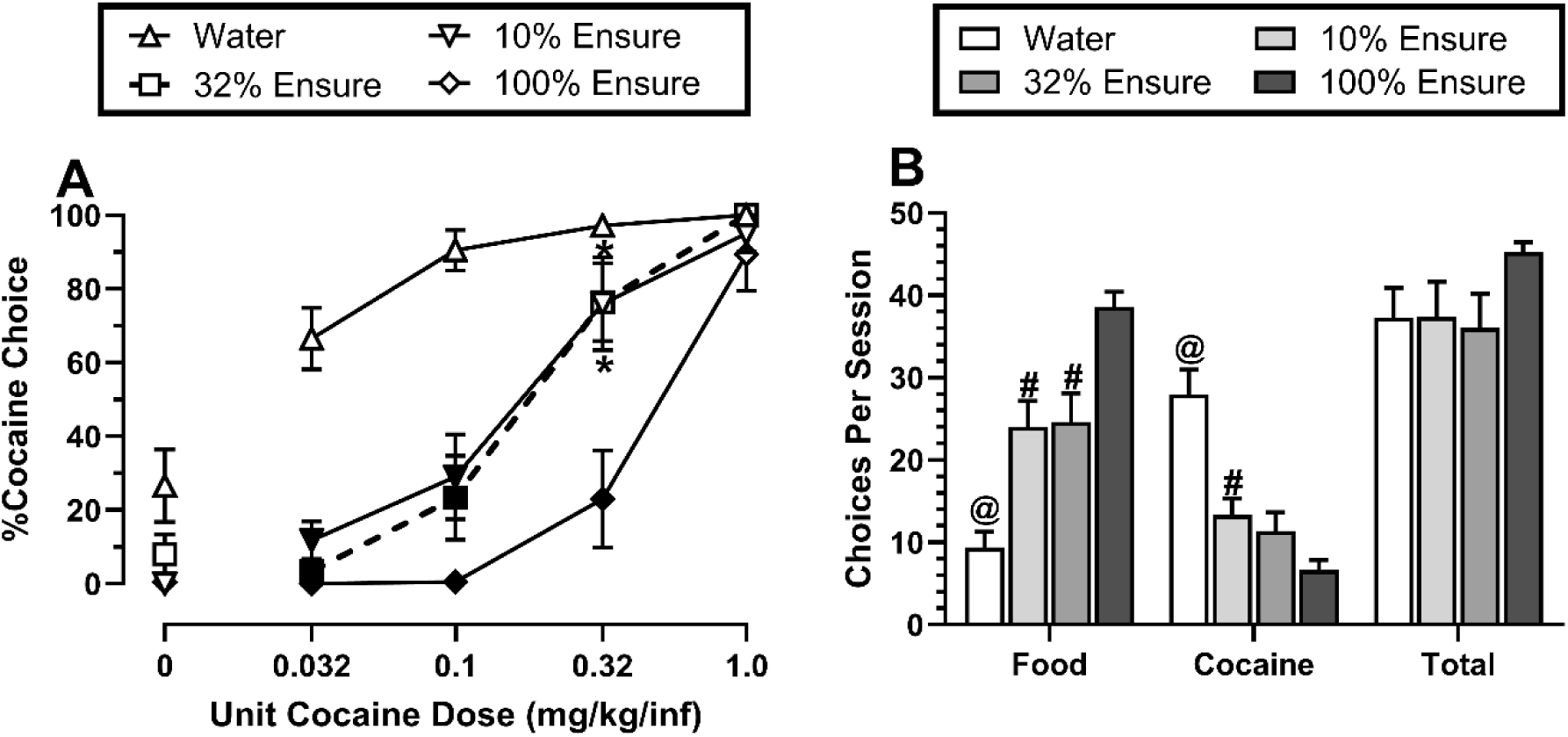
Effects of Ensure concentration on cocaine-vs-food choice in male and female Sprague Dawley rats (N= 7-10, 3-6M/ 4F). A) Percent cocaine choice across various Ensure concentrations. C) Total reinforcers earned per session across various ensure concentrations. All points and bars represent the group means±SEM of the last two days of each 5-day experiment. Filled data points represent a significant (p <0.05) difference compared to water, whereas asterisk represent a significant difference compared to 100% Ensure. ^$^ represents a significant difference between 10% and 100% Ensure. ^@^ represents a significant difference compared to every other ensure concentration, whereas ^#^ represent a significant difference compared to 100% Ensure.

### 3.2 Repeated 5-day Xanomeline treatment on cocaine choice

Figure 2 shows the effects of repeated 5-day vehicle and Xanomeline (1.0 – 10 mg/kg/day) treatment on cocaine-vs-food choice in male and female rats. Figure 2A shows the experimental timeline. Administration of Xanomeline resulted in a significant dose-dependent decrease in %cocaine choice (cocaine dose: F(2.0, 18) = 257, p<0.0001; xanomeline dose: F(2.9, 26.4) = 6.45, p = 0.002; Interaction: F (3.9, 32.6) = 4.76, p = 0.004; Figure 2B), post-hoc analysis showed a significant reduction in %cocaine choice at 5.6 mg/kg (p = 0.01) and 10 mg/kg (p = 0.04). There were no significant reductions in choices per component at any Xanomeline doses tested (Figure 2C). Lastly, there was no significant changes in the total number of reinforcers earned but there does seem to be a downward trend in total cocaine reinforcers and an upward trend in total food reinforcers (Figure 2D).

**Figure 2:**
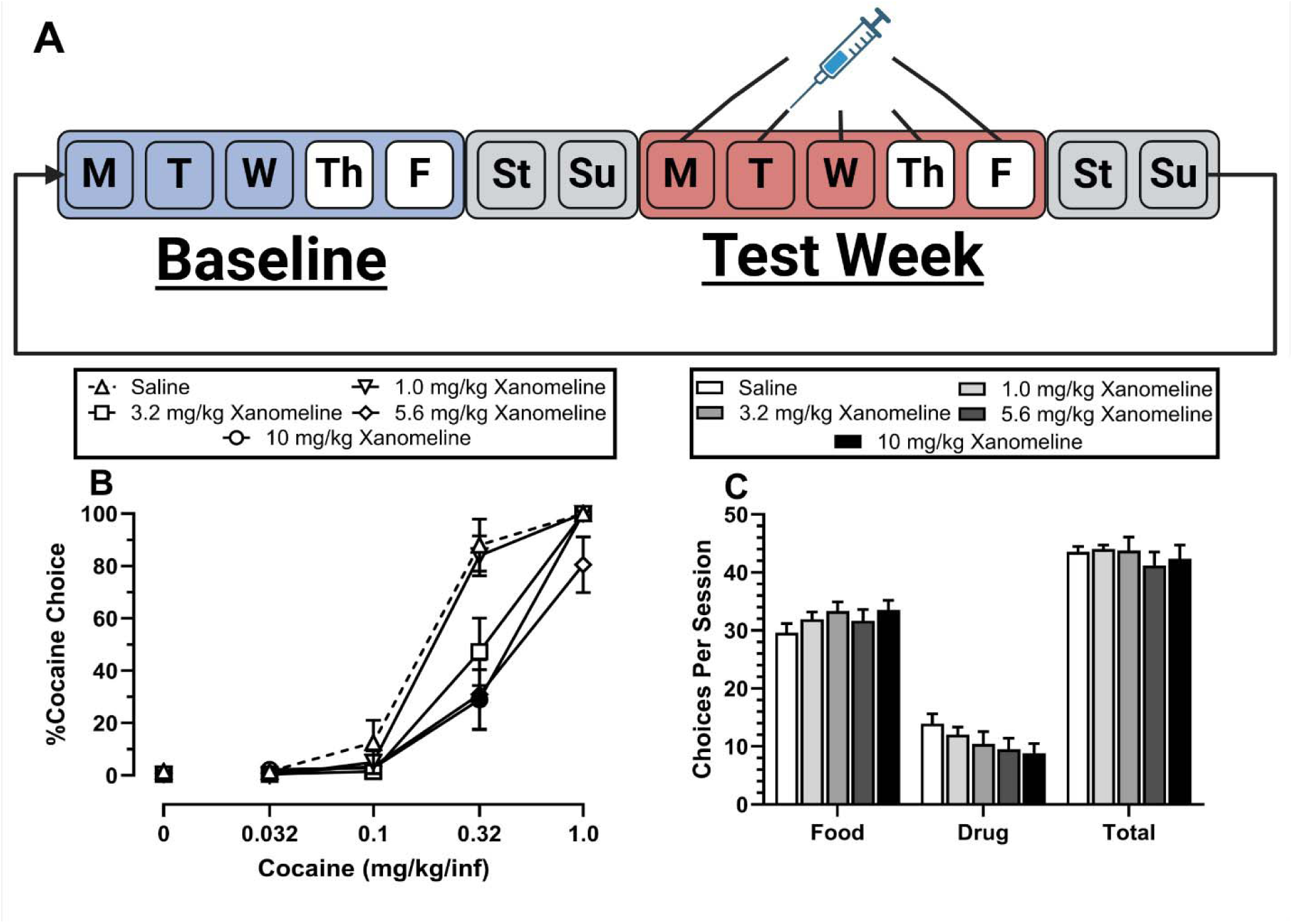
Effects of xanomeline dose on cocaine-vs-food choice in male and female Sprague-Dawley rats (N=9-10, 4M/5-6F). A) Experimental timeline of repeated xanomeline dosing and behavioral assessment. B) Percent cocaine choice during repeated xanomeline administration. C) Total choices earned per session during repeated xanomeline administration. All points and bars represent the average end points during the last two days of weekly testing +SEM. Filled Data points represent a statistically significant difference in percent cocaine choice compared to saline treatment (p<0.05). Figures created with BioRender.com.

### 3.3 Effects of a single VU0364572 administration on cocaine choice

Figure 3 shows the effects of a single vehicle or VU’72 dose administration on cocaine-vs-food choice over two consecutive weeks. Panels 3B,D show vehicle and VU’72 effects on percent cocaine choice, choices per component, and reinforcers per session during the first week. Neither saline nor any VU’72 dose significantly altered any dependent measure (cocaine dose: F (1.2, 11.2)= 81.7, p <0.0001, Figure 3B). Panels 3C, 3E, and 3G show vehicle and VU’72 effects on percent cocaine choice, choices per component, and reinforcers per session during the second week. Figure 3C shows significantly decreased cocaine choice at the 0.32 mg/kg/infusion unit cocaine dose compared to vehicle (cocaine dose: F(1.2, 11.8) = 73.1, p < 0.0001; VU’72 dose: F(0.8, 8.0) = 8.32, p = 0.024; interaction: F(3.2, 16.6) = 3.62, p = 0.033), Post hoc analysis showed a significant decrease at 0.32 mg/kg (p = 0.03). There was no significant effect of VU’72 on choices per component (Figure 3E). However, there was a significant VU’72 × reinforcer interaction (F: 2.1, 9.7) = 7.08, p = 0.012) for Figure 3G. Post-hoc analyses correcting for multiple comparisons failed to detect a significant VU’72 effect on total, cocaine, or food reinforcers earned during the session.

**Figure 3:**
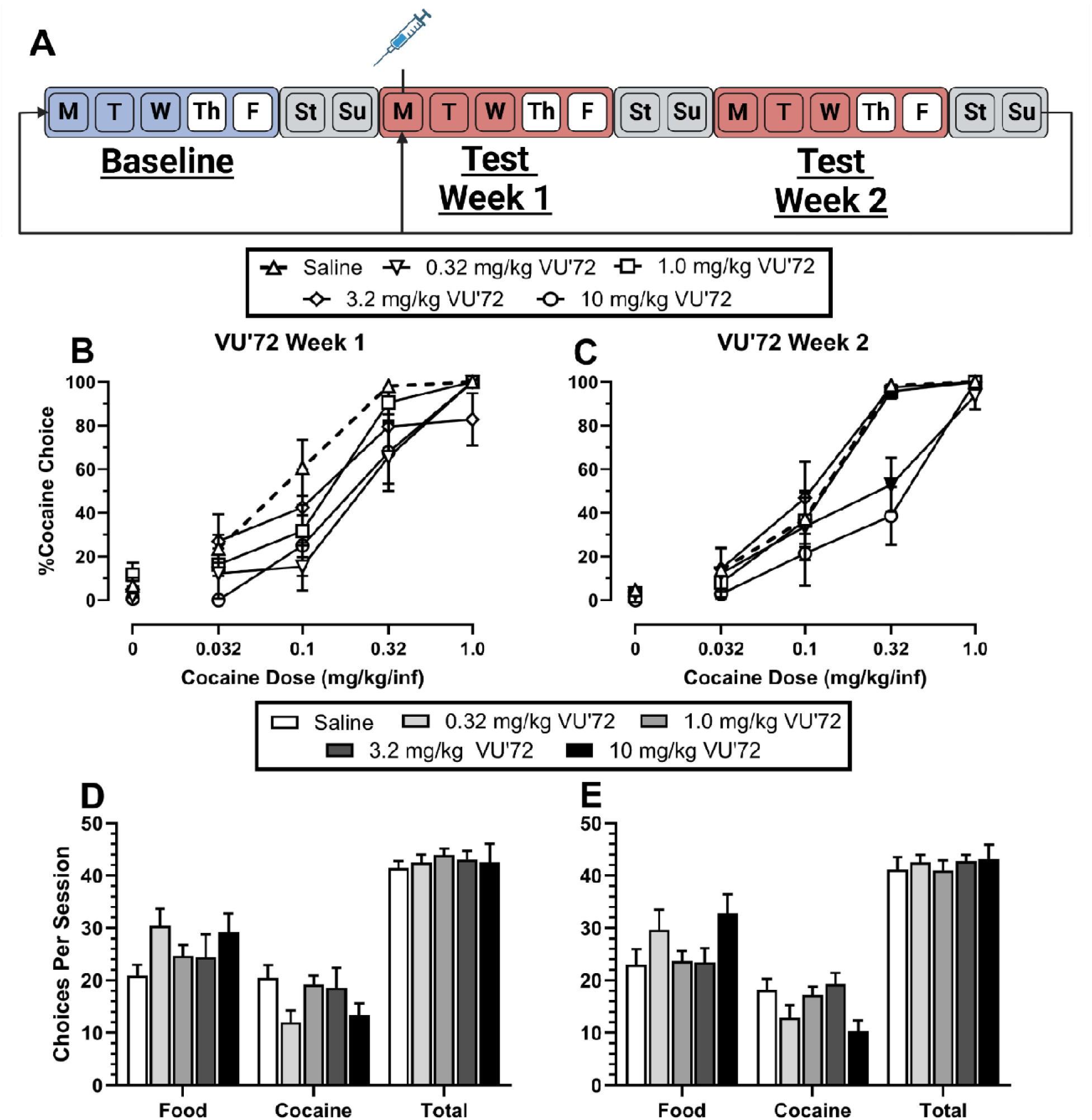
Effects of acute VU0364572 (VU’72) on cocaine-vs-food choice over the course of two weeks in male and female Sprague Dawley rats (N= 7-9, 3-6M/ 2-4F). A) Experimental timeline of acute VU’72 dosing and behavioral assessment. B) Percent cocaine choice during the first week post-acute treatment. C) Percent cocaine choice during the second week post-acute treatment. D) Total choices per session during the first week, post-acute treatment. E) Total choices per session during the second week, post-acute treatment. All points and bars represent the average end points during the last two days of weekly testing +SEM. Filled data points represent a significant difference compared to vehicle (p<0.05). Figures created with BioRender.com.

### 3.4 Repeated 5-day VU0364572 treatment on cocaine choice

Figure 4 shows the effects of repeated 5-day vehicle and VU’72 (0.32 – 10 mg/kg/day) treatment on cocaine-vs-food choice in male and female rats. Figure 4A shows the experimental timeline. There was no significant effect of VU’72 treatment on percent cocaine choice (cocaine dose: F(2.0, 22.3) = 51.7, p<0.0001;Figure 4B), choices per component (Figure 4C), or reinforcers per session (Figure 4D).

**Figure 4:**
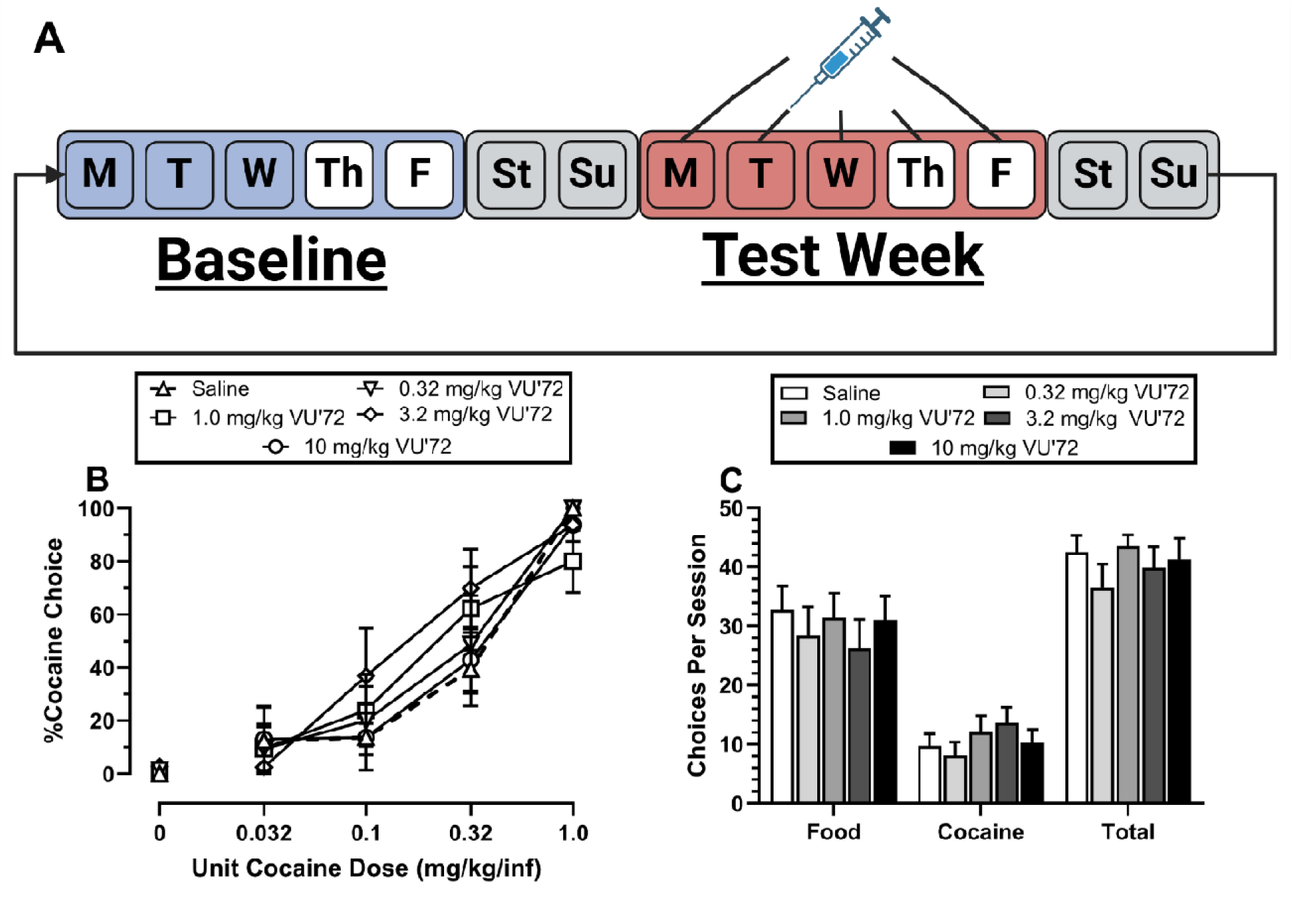
Effects of repeated VU0364572 (VU’72) administration on cocaine-vs-food choice in male and female Sprague Dawley rat (N= 8-9, 3-6M/2-5F). A) Experimental timeline of repeated VU’72 dosing and behavioral assessment. B) Percent cocaine choice during repeated VU’72 administration. C) Total choices earned per session during repeated VU’72 administration. All points and bars represent the average end points during the last two days of weekly testing +SEM. Figures created with BioRender.com.

## 4.0 Discussion

The present study compared the effectiveness of the M1/M4-preferring agonist xanomeline and the M1 bitopic agonist VU0364572 treatments capacity to attenuate cocaine-vs-food choice in male and female rats. There were three main findings. First, repeated 5-day xanomeline treatment significantly attenuated cocaine choice in male and female rats consistent with previously published xanomeline treatment effects in male rats. Second, both acute and repeated 5-day VU’72 treatments failed to meaningfully attenuate cocaine choice in male and female rats. These results are inconsistent with a previous rat study and do not support a robust role of M1 receptors alone in cocaine self-administration. Lastly, increasing the ensure concentration significantly attenuated cocaine choice consistent with the extant literature in rhesus monkeys and rats. Overall, these results support the continued evaluation and development of the M1/M4-preferring agonist xanomeline as a candidate cocaine use disorder pharmacotherapy. Moreover, these results implicate the muscarinic cholinergic system as a potential mechanism in modulating cocaine reinforcement.

Cocaine maintained a dose-dependent increase in choice in both male and female rats, consistent with the extant literature in rats, non-human primates, and humans [25–32]. Furthermore, *increasing* the ensure concentration to 100% attenuated cocaine choice and *decreasing* the ensure concentration facilitated cocaine choice. These results are also congruent with previous cocaine choice experiments in rats and non-human primates [33–37]. Moreover, the present ensure concentration manipulations in this study are consistent with published opioid-vs-food choice studies in rats, suggesting that effects of manipulating alternative reinforcers is consistent across drugs of abuse [25]. These robust reductions in drug choice in response to modifications in alternative reinforcement echo clinical strategies used to treat substance use disorder, specifically contingency management. The focus of contingency management is to promote behavioral reallocation toward alternative reinforcers (i.e. decrease drug choice) by reinforcing behavior that promote abstinence (i.e. increase alternative reinforcer choice) [38]. This non-pharmacological intervention has proven clinically effective as a study showed that subjects undergoing outpatient treatment for cocaine use disorder showed greater prolonged cocaine abstinence with contingency management as comparted to subjects without [39]. Our data shows that our assay produces similar results to contingency management via similar mechanisms (ie. modifying alternative reinforcer strength). Therefore, our ensure manipulations provide a non-pharmacological and clinically relevant positive control in the absence of any FDA approved pharmacological comparators. In conclusion, our concurrence with the extant clinical and pre-clinical literature shows that our assay is built on an empirical foundation that supports the exploration of novel pharmacotherapies.

Repeated 5-day xanomeline treatment, dose-dependently decreased cocaine choice in male and female Sprague-Dawley rats, indicating that xanomeline treatment reduced the relative reinforcing strength of cocaine. The present data, that 5.6 and 10 mg/kg/day xanomeline treatment attenuated cocaine choice, were consistent with a previously published study in male rats [22]. We expand upon this previous finding by including females, further demonstrating that this reduction in cocaine self-administration does not appear to be sex specific. Moreover, the present results are consistent with previous data showing that xanomeline decreased sessions required to meet extinction criteria and cocaine induced reinstatement in mice [40]. Additionally, repeated xanomeline treatment decreased cocaine choice with similar effectiveness to increasing the magnitude of the alternative nondrug reinforcer further supporting the potential of xanomeline as a candidate cocaine use disorder pharmacotherapy.

In contrast to Xanomeline, VU0364572 treatment effects on cocaine choice were inconsistent with previous data during acute or repeated dosing. There was a significant decrease in cocaine choice with 0.32 and 10 mg/kg VU’72 during the second week post-acute dosing; however, these results were not reproduced during repeated dosing nor these doses significant compared to a consolidated baseline (data not shown). The present results are in contrast to a previous study that reported a single 1.0 mg/kg VU’72 administration resulted in a prolonged attenuation of cocaine choice in male Sprague-Dawley rats [20]. Examination of VU’72 treatment effects on cocaine choice by sex did not unmask a VU’72 treatment effect in either sex. Therefore, the present data does not support the hypothesis that selective activation of M1 receptors robustly modulates cocaine self-administration and that VU’72 would be a viable cocaine use disorder pharmacotherapy. This brings into question why xanomeline decreased cocaine choice while VU’72 failed to.

The primary difference between xanomeline and VU’72 is their activity on muscarinic receptors, as xanomeline functions as a M1/M4 preferring muscarinic agonist and VU’72 acts soley as an M1 allosteric agonist. Thus, we have two possible hypotheses to explain the differential effects between our ligands:

The first is that the reduction in cocaine reinforcement is primarily derived from M4 muscarinic receptor activation. There is sufficient neurobiological evidence to suggest that M4 receptor co-expresses with D1 receptor within the nucleus accumbens, meaning that activation of M4 receptors would have an antagonist relationship to D1 activation within the direct dopamine pathway [15,16]. This antagonism to D1 receptor activation could be the primary reason for the reductions in the reinforcing effects of cocaine [41,42]. There is also the possibility that M4 receptor activation is causing a reduction in dopamine release as well. However, previous experimental data shows administration of xanomeline causes an increase in dopamine release within the NAc and mPFC [43]. Therefore, the reductions in cocaine choice are more likely to be attributed to reductions in post-synaptic D1 signaling than reductions in dopamine release within the synapse itself. There is also a wealth of preclinical behavioral experiments that suggest that M4 activation alone reduces cocaine reinforcement. Previous experiments in mice have shown that knockout of the M4 muscarinic receptors produces a significant increase cocaine self-administration, progressive ratio breakpoint, and cocaine induced dopamine efflux compared to wildtype [44]. Additionally, behavioral experiments in rats using the M4 PAM VU0152100 showed a significant, behaviorally selective, and dose dependent decrease in cocaine self-administration as well as reductions in cocaine induced dopamine release within the striatum [45]. Furthermore, treatment with the M4 PAM VU0152099 induced a significant attenuation in cocaine choice during repeated treatment similar to our xanomeline data [46]. Therefore, there is neurological and behavioral evidence that the decreases in cocaine reinforcement may be due solely to activation of the M4 muscarinic receptor.

The second hypothesis is that activation of both M1 and M4 muscarinic receptors is necessary to blunt cocaine reinforcement. This hypothesis still relies on the previous evidence of M4 muscarinic receptor activity blunting cocaine reinforcement. However, it proposes that the M1 muscarinic receptors expressed on glutamatergic projections to the NAc from the PFC, inhibiting D2 receptor activation in the indirect dopamine pathway [17,18]. This is further supported by experimental evidence, using whole-cell patch clamp electrophysiology, that M1 muscarinic receptors modulate excitatory inputs into the NAc [18]. Therefore, xanomeline could be blunting both the direct and indirect dopamine signals in the NAc to reduce cocaine’s reinforcing effects. This would also be congruent with recent behavioral experiments that showed that both M1 and M4 muscarinic receptors were necessary to achieve maximal blockade of cocaine’s discriminative stimuli in a drug discrimination assay [47]. Additional evidence showed a behaviorally selective reduction in IV cocaine self-administration acquisition in mice only during a high dose of <u>both</u> VU0364572 (M1 PAM) and VU0152100 (M4 PAM), without any significant decreases in acquisition in either group when administered alone [48]. This may also explain why we saw significant effects with xanomeline but not in our VU’72 data, as we have two effects working in concert to produce a more robust attenuation of cocaine choice than VU’72 can produce alone. Regardless of the mechanism by which xanomeline is attenuating cocaine choice, the future directions of our experiment are clear.

To answer the presented hypothesis, further experiments need to be run that look at the effect of M4 muscarinic receptor activation on cocaine choice. By utilizing the selective M4 PAM VU0152100, we should be able to decipher the effects of M4 activation on the reinforcing effects of cocaine. If there is significant attenuation that would be further evidence to support that M4 is the primary mediator of the reduction in cocaine’s reinforcing effects. However, if there is no significant attenuation then there would be further evidence that both M1 and M4 muscarinic receptor activation is required for behaviorally relevant reductions in cocaine’s reinforcing effects.

Regardless of future results, xanomeline functioned to dose-dependently reduce cocaine choice and total reinforcer data trends toward behavioral reallocation without any significant rate decreasing effects. With xanomelines recent approval as an FDA pharmacotherapy, further efforts should be made to explore its utility to treat cocaine use disorder clinically. Additionally, further pre-clinical work should be done to explore the exact interplay of M1 and M4 muscarinic receptors on the mesocortical dopamine pathways.

